# Avatar facial expressions enhance social presence and inter-brain synchronization in virtual reality

**DOI:** 10.64898/2026.01.28.702209

**Authors:** Haruki Kodama, Katsuki Higo, Sotaro Shimada

## Abstract

Virtual reality (VR) has become a prominent medium for computer-mediated social interaction, yet the psychological and neural mechanisms underpinning avatar-mediated communication remain insufficiently understood. This study investigated how dynamically modulated avatar facial expressions shape interpersonal interaction in VR. Pairs of participants engaged in a collaborative creativity task while interacting via avatars whose facial expressions were either amplified, natural, or expressionless. We collected subjective measures of body ownership, social presence, and interpersonal attraction, while simultaneously recording neural activity using functional near-infrared-spectroscopy-based hyperscanning to assess inter-brain synchrony (IBS). Avatars displaying facial expressions, particularly amplified ones, significantly enhanced body ownership, social presence, and interpersonal attraction compared to expressionless avatars. At the neural level, visible avatar facial expressions were associated with IBS in the right temporo-parietal junction (TPJ) and the dorsolateral prefrontal cortex (dlPFC), regions that are commonly implicated in social-cognitive processes such as mentalizing and executive control. Although task performance did not differ across conditions, social presence was positively correlated with creative performance, suggesting that psychological connectedness supports collaborative creativity. Together, these findings indicate that avatar facial expressivity functions as a critical nonverbal social cue that facilitates affective and cognitive alignment between interaction partners. By linking subjective experience with inter-brain neural dynamics, this study provides empirical guidance for the design of virtual environments that promote social engagement and effective collaboration.

## 1. Introduction

Computer-mediated communication has become an integral part of everyday life, encompassing social networking, collaborative work, and education. Among these technologies, virtual reality (VR) is a compelling medium, allowing users to share immersive spaces and interact through embodied digital representations—avatars—that can mimic gestures, gaze, and facial expressions (Gunkel et al., 2018; Seymour et al., 2021). Compared with traditional video conferencing or text-based communication, VR affords a richer channel for nonverbal interaction, allowing subtle bodily and expressive cues to be dynamically integrated into social exchange. Despite the rapid growth of social VR platforms, however, the psychological and neural mechanisms through which users establish and maintain interpersonal connections in such mediated environments remain insufficiently understood. Elucidating how people establish psychological connections in mediated environments is essential for designing technologies that effectively support communication and collaboration.

### 1.1. Facial expressions and social presence in mediated environments

Facial expressions are among the most critical nonverbal cues in interpersonal communication. They signal internal states, such as emotions and intentions, and regulate social relationships by inviting or discouraging approach behaviors (Krumhuber et al., 2013; Reed et al., 2014). In physical face-to-face communication, expressive facial cues enhance perceived empathy, trustworthiness, and affiliation (Seidel et al., 2010; van’t Wout & Sanfey, 2008). When these cues are absent or ambiguous, as is common in mediated settings, communication can become emotionally flat and cognitively effortful.

According to Social Presence Theory (Short et al., 1976), the sense of “being with” another person in a mediated environment depends on the richness and immediacy of social cues. Subsequent models conceptualized social presence as a multidimensional construct involving psychological involvement, co-presence, and behavioral engagement (Biocca et al., 2003). From this perspective, avatars capable of reproducing realistic facial expressions are not mere visual placeholders but are socially responsive entities that can elicit empathic and affiliative responses similar to those of real partners. Empirical studies have confirmed that higher avatar expressivity increases perceived intimacy and rapport (Bente et al., 2008; Oh et al., 2018; Oh Kruzic et al., 2020; Kyrlitsias & Michael-Grigoriou, 2022). Therefore, enhancing the nonverbal capabilities of avatars may be a key strategy for achieving authentic social experiences in VR.

While Social Presence Theory emphasizes the perception of others, the Proteus effect (Yee & Bailenson, 2007: Sah et al., 2021; Coesel et al., 2024; Liu, 2023) highlights changes in self-perception. Avatars shape not only how others perceive us but also how we perceive ourselves. For instance, individuals embodied in attractive or dominant avatars tend to act more confidently and exhibit greater social engagement (Yee & Bailenson, 2007). This phenomenon demonstrates that digital embodiment is not a neutral interface but an interactive psychological process linking perception, action, and self-representation. Recent comprehensive reviews further consolidate the Proteus effect as one of the canonical findings in virtual reality research, highlighting that self-avatars systematically shape user behavior across perceptual, cognitive, and social domains (Bailenson et al., 2025). Collectively, these theories describe the dual nature of avatar-mediated embodiment: how we perceive others and how we act through our virtual selves.

Recent work has extended the Proteus effect to dyadic and group contexts, demonstrating that expressive, human-like avatars can enhance mutual understanding, empathy, and cooperation (Dubosc et al., 2021; Roth et al., 2018). These findings suggest avatar design influences both intrapersonal (self-related) and interpersonal (relationship-related) dimensions of virtual experiences. However, most prior studies have relied on static or pre-programmed facial expressions. The advent of VR facial tracking has enabled the projection of real-time dynamic expressions—including subtle eye movements, blinking, and mouth movements—onto avatars. This has opened up new possibilities for studying how dynamic cues affect users’ psychological connectedness.

Beyond simply reproducing natural expressions, recent work has begun to consider whether enhancing or exaggerating facial movements could improve communication in virtual environments— a concept that we refer to here as facial expression amplification. Exaggerating subtle movements (e.g., smiles, eye movements) may make avatars more readable and socially engaging, particularly under conditions of limited resolution or network latency. However, the psychological effects of amplified expressions remain largely unexplored. Clarifying whether such amplification enhances connectedness or undermines authenticity is essential for understanding how emotional cues function in technologically mediated social interactions.

## 1.2. Neural mechanisms of social interaction: inter-brain synchrony (IBS)

In parallel with behavioral studies, social neuroscience has begun investigating how the brain coordinates during real-time interaction. Inter-brain synchrony (IBS)—the temporal alignment of neural activity between individuals—has emerged as a robust neural marker for cooperation and collaborative behavior (Cui et al., 2012; Dikker et al., 2014; Redcay & Schilbach, 2019; Shiraishi & Shimada, 2021; Ogawa & Shimada, 2023). Techniques such as functional near-infrared-spectroscopy (fNIRS) hyperscanning permit the measurement of this coupling in naturalistic settings while participants communicate or perform tasks jointly (Cui et al., 2012; Funane et al., 2011; Koide & Shimada, 2018). Increased synchrony within regions of the social brain network, particularly the temporoparietal junction (TPJ), has been linked to higher levels of mutual understanding and cooperative behavior, reflecting its central role in perspective-taking and mentalizing (Lu et al., 2019; Tang et al., 2016). Thus, interpersonal neural coupling provides an objective window into the depth of social connectedness, complementing subjective questionnaire measures.

Applying this framework to VR interactions is a relatively new but promising endeavor. If expressive avatars facilitate perspective-taking and emotional contagion, they should, in turn, enhance neural synchrony between users, indicating deeper cognitive alignment and shared intentionality. Measuring these effects could reveal how technological mediation shapes fundamental social processes at both psychological and neural levels.

Although increasing evidence from human–computer interaction, psychology, and neuroscience has highlighted the importance of social connectedness in mediated communication, these perspectives have not yet been fully integrated. Research on VR communication has often prioritized user experience or interface design, whereas cognitive and neural mechanisms underlying social interaction remain less explored. Conversely, social neuroscience research has largely relied on highly controlled experimental paradigms and single-brain measurements, which limit the investigation of dynamic and reciprocal social interactions in ecologically valid settings (Koike et al., 2015). As a result, commonly used tasks such as gaze-following or joint attention paradigms may be insufficient for capturing the complexity of social interaction in immersive virtual environments. Bridging these domains would advance theory and inform the design of socially intelligent systems that can sense and adapt to users’ affective states. The present study addresses this gap by combining systematic manipulations of behavioral realism in VR with fNIRS-based hyperscanning, enabling the simultaneous assessment of subjective, behavioral, and neural indicators of social connectedness. This integrative approach is theoretically grounded in the view that social presence arises from embodied coupling between interacting individuals. When avatars reproduce realistic facial cues, they may elicit stronger sensorimotor resonance and emotional attunement, leading to higher perceived presence and attraction. Such experiences are expected to manifest as increased inter-brain synchrony in social-cognitive regions.

### 1.3. Aims of the current study

Building on these theoretical considerations, this study investigated how dynamically modulated avatar facial expressions influence interpersonal experiences and neural synchronization during virtual interaction. We manipulated avatar expressivity across three conditions (Amplified, Natural, and None) using real-time facial tracking that mapped participant expressions onto avatars with different intensity weights. Participant pairs engaged in a collaborative creativity task while their brain activity was simultaneously recorded using fNIRS hyperscanning. Subjective measures of body ownership, social presence, and interpersonal attraction were collected via questionnaires after each condition.

This multimodal design allowed us to examine how variations in avatar expressivity affect both subjective experience and objective neural coupling. Specifically, we tested the following hypotheses: (1) Avatars displaying facial expressions—particularly amplified ones—enhance body ownership, social presence, and interpersonal attraction compared to expressionless avatars. (2) These enhanced subjective experiences are accompanied by greater IBS in regions associated with social cognition (e.g., the TPJ). (3) Social presence positively correlates with collaborative creativity, indicating that psychological connectedness supports joint performance. By integrating behavioral and neural evidence, this study offers a more comprehensive understanding of how expressive avatars mediate human behavior in virtual environments.

## 2. Materials and methods

### 2.1. Participants

A total of 32 healthy participants (16 same-gender pairs; 3 female and 13 male pairs; mean age = 21.65 ± 0.54 years) participated in the experiment. Data from 15 pairs were used for the fNIRS analysis; one pair was excluded owing to measurement issues. Before the experiment, all participants were provided with an overview of the study and safety information, and subsequently provided signed informed consent.

### 2.2. Experimental environment and equipment

Participants were seated face-to-face, each wearing a head-mounted display and facial tracker (Vive Pro Eye + Vive Facial Tracker, HTC, Taipei, Taiwan) and holding handheld controllers. This physical arrangement was mirrored in the VR environment, where participants also sat face-to-face. The Vive Pro Eye tracked gaze, the Vive Facial Tracker captured mouth movements, and the controllers recorded arm movements; these inputs were reflected in real-time on their avatars. The virtual environment was developed using the Unity game engine (version 2019.4.31f1, Unity Technologies, San Francisco, CA, USA).

Two male and two female avatar models were created using VRoid Studio (Pixiv Inc., Tokyo, Japan). Each model was used across three facial expression conditions: (1) Amplified (expression weights set to 200%); (2) Natural (expression weights set to 100%); and (3) None (expression weights set to 0%) (Fig.1). Specifically, avatars were designed with predefined facial shapes using BlendShape, a method for controlling facial expressions. These expressions were generated in real-time based on data from the facial tracker. In the Amplified condition, the avatar’s BlendShape values were set to twice those in the Natural condition, meaning any expression change was visually amplified by a factor of two. In the None condition, no facial expression data was applied to the avatar, rendering it expressionless.

**Fig. 1.**
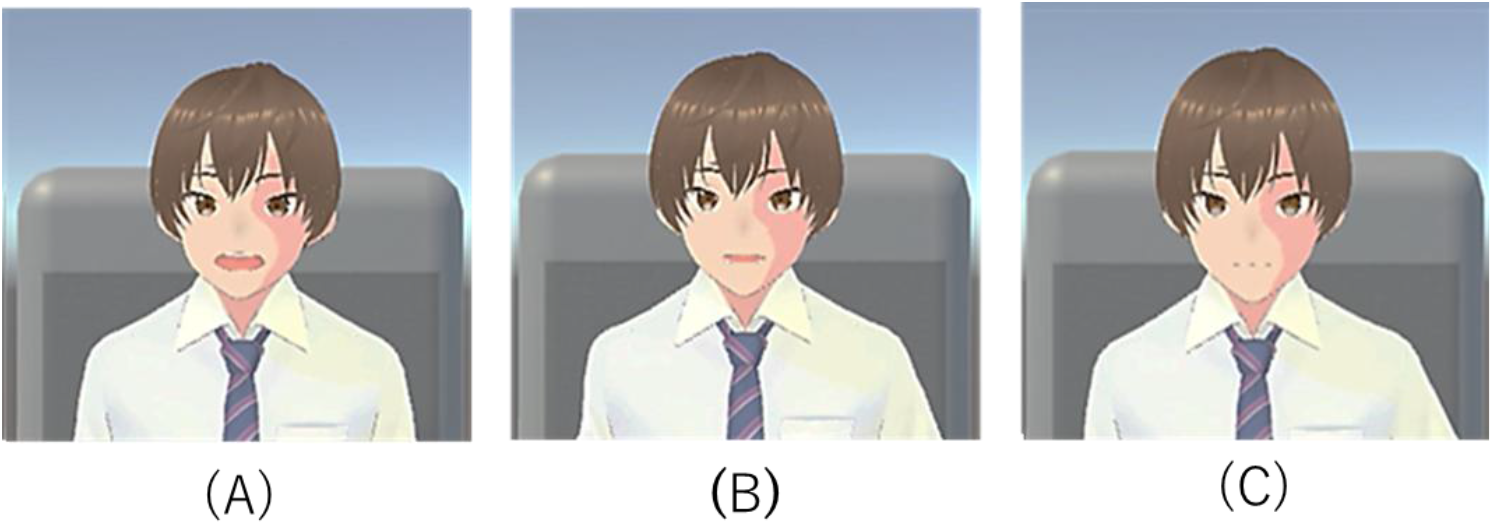
One of the avatar models used in the experiment, displaying the three expression conditions: (A) Amplified, (B) Natural, and (C) None.

**Fig. 2.**
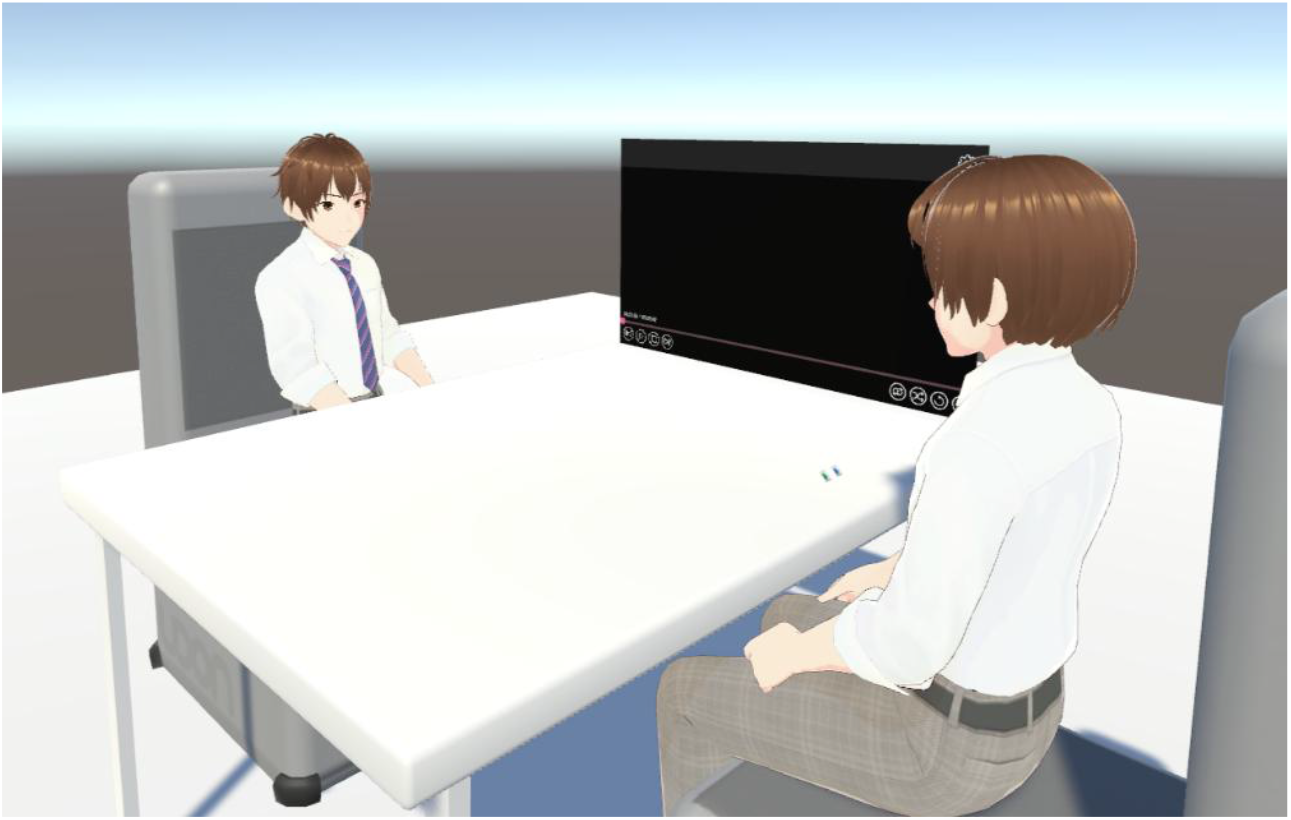
Experimental setup showing the interpersonal interaction scene within the virtual reality (VR) environment.

### 2.3. Experiment

Participants in each pair were seated facing one another. The procedure comprised three instruction blocks, each followed by a 5-min task block. After completing each task block, participants filled out a questionnaire. The three task blocks corresponded to the three avatar conditions (Amplified, Natural, and None), and the order of conditions was counterbalanced across participant pairs. The interpersonal interaction task was a collaborative version of the Alternative Uses Task (AUT). Participants were instructed to collaboratively generate as many alternative uses as possible for common everyday objects. Task performance was assessed using two standard metrics: fluency and originality (Guilford, 1967; Okuda et al., 1991). Fluency was defined as the total number of ideas generated by the pair (i.e., the sum of both individuals’ idea counts). Originality was evaluated using a subjective rating procedure.

Two independent raters assessed each idea on a five-point Likert scale from 1 (“not original at all”) to 5 (“highly original”). Inter-rater reliability, assessed using the intraclass correlation coefficient, was 0.66. The final originality score for each idea was computed as the average of the two raters’ ratings.

### 2.4. Questionnaire items

The post-task questionnaire assessed three constructs: body ownership, social presence, and interpersonal attraction. All items were rated on a seven-point Likert scale ranging from –3 (“strongly disagree”) to +3 (“strongly agree”).

Body Ownership, defined as the sense that avatar’s body parts belong to oneself, was measured using items adapted from the self-body recognition questionnaire used by Kalckert and Ehrsson (2012). As this study did not involve significant body movement, items related to the sense of agency were excluded. Social Presence, or the “sense of being with others,” depends on the ease with which one can recognize “the intelligence, intentions, and sensory impressions of others” (Biocca, 1997). It was measured with items adapted from the Social Presence Questionnaire (Harms and Biocca, 2004). Interpersonal attraction, i.e., the positive or negative attitude one holds toward another person, was measured using the items employed by Oh Kruzic et al. (2020).

Questionnaire scores for body ownership, social presence, and interpersonal attraction were compared across the three experimental conditions (Amplified, Natural, and None). The Shapiro–Wilk test indicated the data were not normally distributed. Consequently, a Friedman test was performed for statistical analysis. Post-hoc multiple comparisons were conducted using the Wilcoxon signed-rank test, with p-values adjusted via Holm’s method.

### 2.5. fNIRS measurement

Hemodynamic responses, specifically changes in oxygenated hemoglobin (oxy-Hb) concentrations, were assessed using a 48-channel (ch.) fNIRS system (OMM-3000; Shimadzu Corporation), with 24 channels allocated to each participant. Based on prior research highlighting the involvement of the temporoparietal junction (TPJ) in social interaction and group creativity (Cheng et al., 2015; Decety, 2010; Funane et al., 2011; Goel et al., 2015; Sun et al., 2016; Tang et al., 2016), we selected a measurement region encompassing the TPJ in each participant (Fig. 3). Optode locations were confirmed in Montreal Neurological Institute (MNI) space using the probabilistic estimation method with a 3D digitizer (FASTRAK, Polhemus, Colchester, VT, USA) (Singh & Okamoto, 2005).

**Fig. 3.**
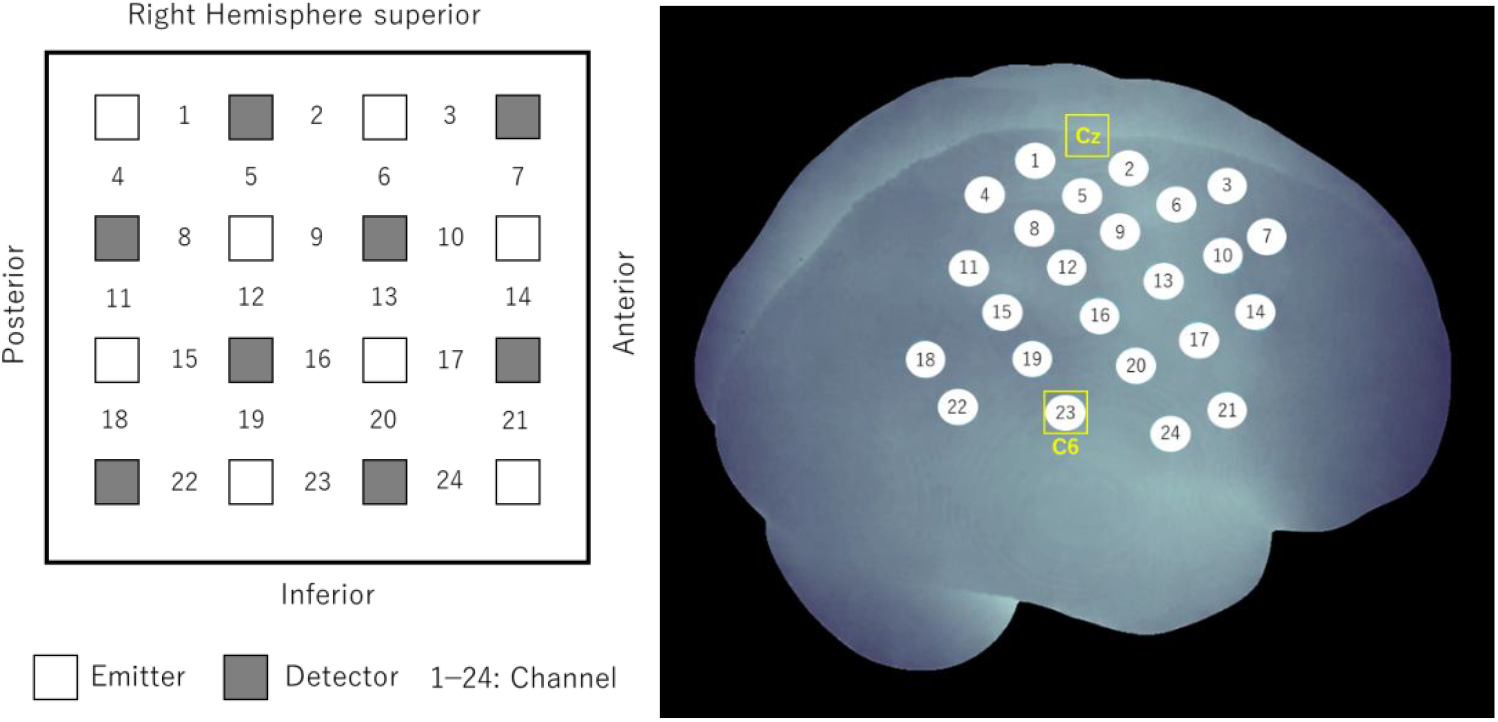
Schematic diagram of the fNIRS optode placement (24 channels) on the right hemisphere of each participant. C6 was approximately located at channel 23. The optodes of the bottom line were placed as horizontally as possible.

To quantify inter-brain synchrony (IBS), we employed a general linear model (GLM)-based fitting approach (Hasson et al., 2004). To this end, IBS was computed not only for actual interacting pairs but also for randomly paired participants, allowing us to examine whether observed neural synchrony was attributable to genuine social interaction rather than shared task structure or common sensory input. In this framework, the neural activity time series of one participant was used as the design matrix to model the simultaneously recorded neural activity of their interaction partner. Specifically, for each measurement channel, the time course from a given participant was used to predict the activity of the corresponding channel at the same scalp location in the partner, yielding a channel-wise index of IBS. This analysis was performed using the neural activity time series recorded during the task periods. As the resulting IBS values deviated from normality, as confirmed by the Shapiro–Wilk test, statistical evaluation was performed using an aligned rank transform analysis of variance (ART-ANOVA). To control for multiple comparisons across channels, false discovery rate (FDR) correction was applied.

## 3. Results

### 3.1. Questionnaire

The questionnaire results for Body Ownership, Social Presence, and Interpersonal Attraction are presented in Fig. 4. A Friedman test was conducted to analyze the main effect of facial expression (Amplified, Natural, None) on these scores. A significant main effect of facial expression was found for Body Ownership (χ^2^ (2) = 7.06, *p* < 0.05 ; Fig. 4A). Post-hoc Wilcoxon signed-rank tests revealed that the Amplified condition resulted in significantly higher body ownership scores than the None condition (*p* < 0.05, *z* = 2.54, *r* = 0.457). A significant main effect was also observed for Social Presence (χ^2^ (2) = 16.3, *p* < 0.001; Fig. 4B). Post-hoc analysis showed that the Amplified condition yielded significantly higher social presence scores than both the Natural (*p* < 0.01, *z* = 3.02, *r* = 0.542) and None (*p* < 0.01, *z* = 3.55, *r* = 0.637) conditions. Finally, a significant main effect of facial expression was found for Interpersonal Attraction (χ^2^ (2) = 7.83, *p* < 0.05; Fig. 4C). Post-hoc analysis revealed that both the Amplified (*p* < 0.05, *z* = 2.52, *r* = 0.452) and Natural (*p* < 2.53, *z* = 2.53, *r* = 0.455) conditions resulted in significantly higher interpersonal attraction scores than the None conditions.

**Fig. 4.**
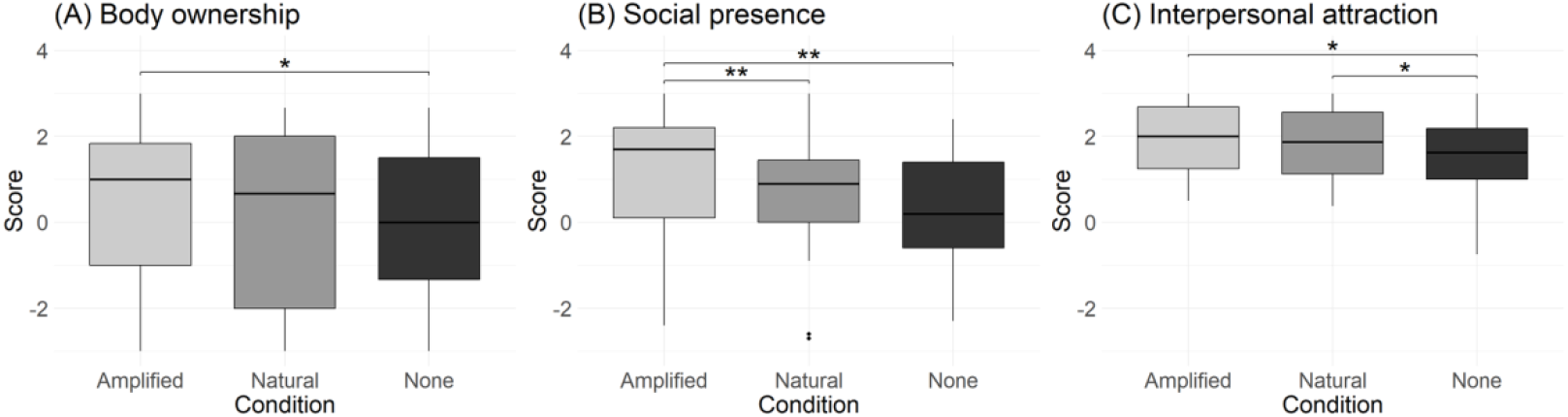
Questionnaire results for the three experimental conditions. Boxplots show scores for (A) Body ownership, (B) Social presence, and (C) Interpersonal attraction. For all plots, significance is indicated as \**p* < 0.05 and \*\**p* < 0.01.

### 3.2. Task scores

The effects of the avatar condition (Amplified, Natural, None) on creativity were examined by analyzing fluency and originality scores. The Friedman test revealed no significant main effect of condition on fluency scores (χ^2^ (2) = 0.036, *p* = 0.982). Mean fluency scores were comparable across conditions: Amplified condition (mean = 17.8, SD = 6.80), Natural condition (mean = 18.0, SD = 6.21), and None condition (mean = 17.9, SD = 6.09). A one-way repeated-measures analysis of variance (ANOVA) indicated no significant main effect of condition on originality scores (*F*(1.67, 25.10) = 0.15, *p* = 0.83, *η*^2^ = 0.15)). Mean originality ratings were similar across conditions: Amplified condition (mean = 2.96, SD = 0.310), Natural condition (mean = 2.96, SD = 0.311), and None condition (mean = 2.94, SD = 0.377).

### 3.3. fNIRS

A 2 (Pair Type: actual vs. random) × 3 (Avatar Condition: Amplified, Natural, None) ART-ANOVA was performed on the computed IBS values revealed a significant interaction in the Ch 7 (Fig.5) and Ch 18 (Fig.6). Probabilistic anatomical estimation based on MNI space using a 3D digitizer indicated that Ch 7 was localized to the middle frontal gyrus, corresponding to the dorsolateral prefrontal cortex (dlPFC), whereas Ch 18 was localized to the angular gyrus within the temporoparietal junction (TPJ). Subsequent analyses therefore refer to these regions as the dlPFC and TPJ, respectively.

**Fig. 5.**
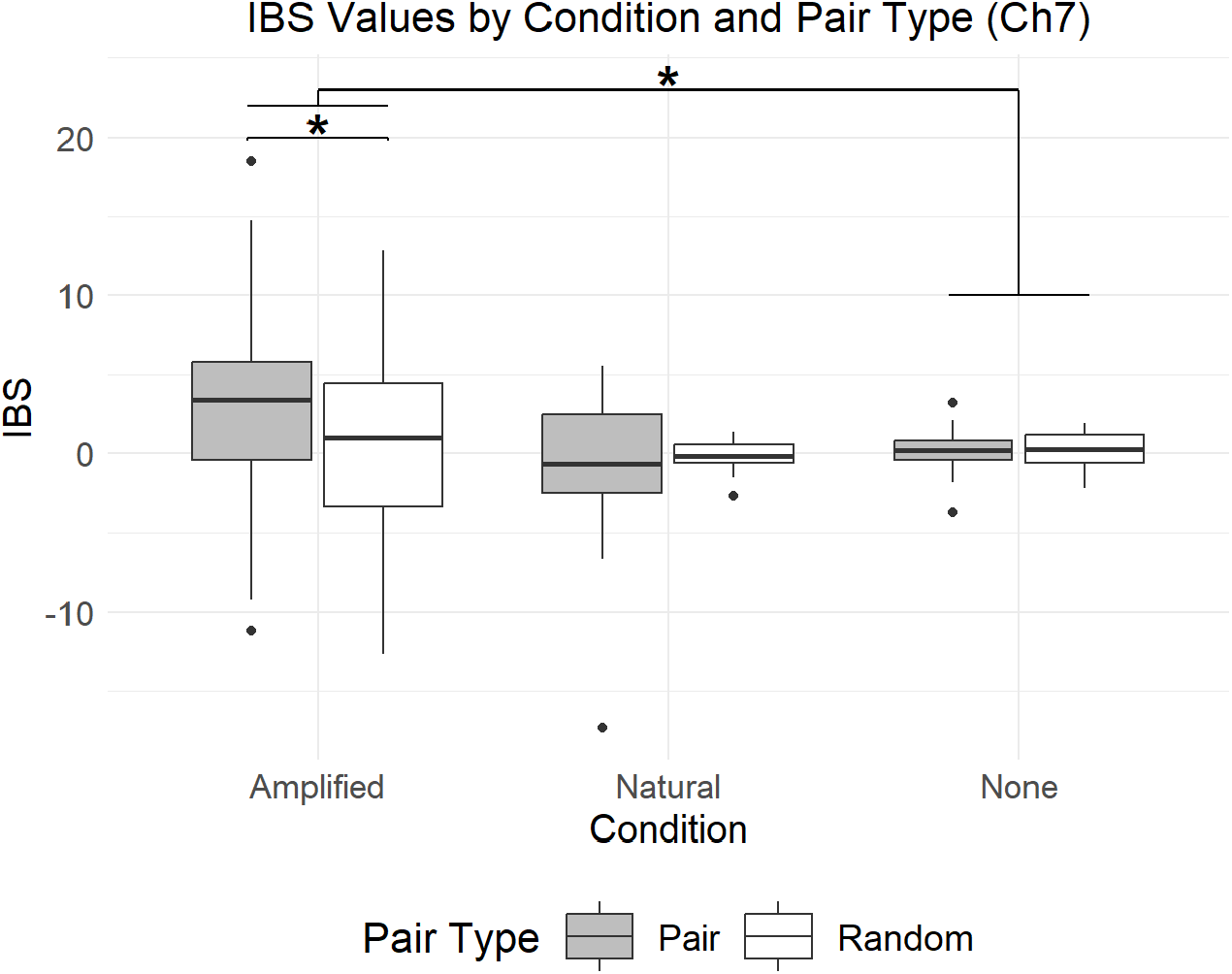
Comparison of inter-brain synchrony (IBS) values across conditions and pair types (Ch 7). IBS was significantly higher for actual interacting pairs (gray bars) than for random pairs (white bars) in the Amplified and Natural conditions. Significance is indicated as \**p* < 0.05.

**Fig. 6.**
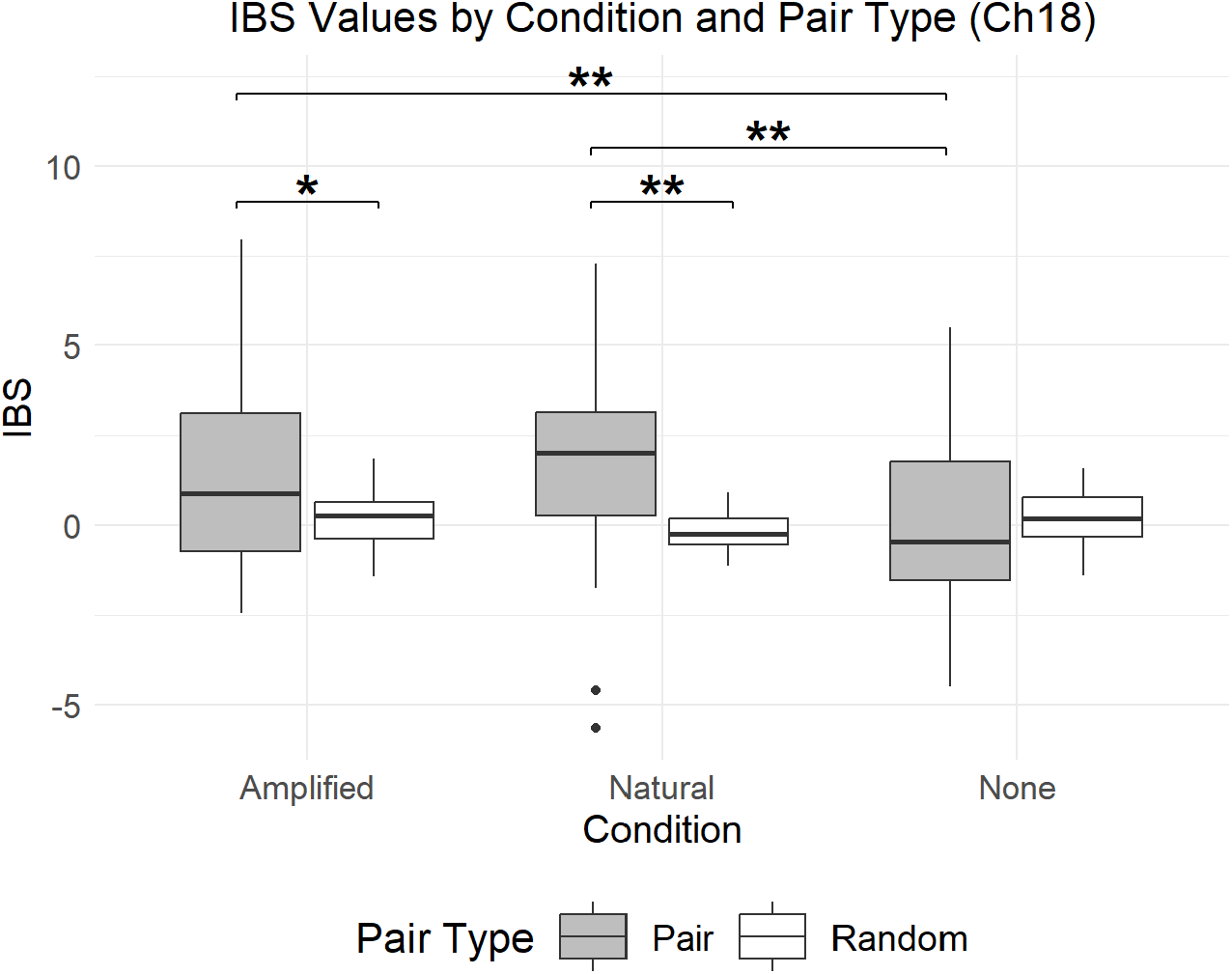
Comparison of inter-brain synchrony (IBS) values across conditions and pair types (Ch 18). IBS was significantly higher for actual interacting pairs (gray bars) than for random pairs (white bars) in the Amplified and Natural conditions. Significance is indicated as \**p* < 0.05 and \*\**p* < 0.01.

In the dlPFC, a significant main effect of Avatar Condition was observed (F(1,58) = 4.33, p < 0.05, η_p^2^ = 0.130), along with a significant interaction between Pair Type and Avatar Condition (F(1,58) = 7.25, p < 0.01, η_p^2^ = 0.200). Post-hoc Wilcoxon signed-rank tests comparing actual and random pairs within each avatar condition revealed that, in the Amplified condition, actual pairs exhibited significantly higher IBS than random pairs (p < 0.05, z = 2.20, r = 0.284).

In the TPJ, significant main effects were found for both Pair Type (F(1,29) = 5.75, p < 0.05, η_p^2^ = 0.165) and Avatar Condition (F(2,58) = 6.48, p < 0.01, η_p^2^ = 0.183), as well as a significant interaction between the two factors (F(2,58) = 16.10, p < 0.001, η_p^2^ = 0.357). Post-hoc Wilcoxon signed-rank tests indicated that, within the actual-pair condition, both the Amplified (p < 0.01, z = 2.65, r = 0.342) and Natural (p < 0.01, z = 2.65, r = 0.342) avatar conditions elicited significantly higher IBS than the None condition. In contrast, no significant differences among avatar conditions were observed in the random-pair condition (χ^2^(2) = 0.467, p = 0.792).

Further comparisons between actual and random pairs within each avatar condition confirmed that actual pairs exhibited significantly higher IBS in the Amplified (p < 0.05, z = 2.18, r = 0.281) and Natural (p < 0.01, z = 3.50, r = 0.452) conditions, indicating that these increases were attributable to genuine social interaction rather than shared task structure.

### 3.4. Correlation analysis

To examine the relationships among the subjective, neural, and performance measures, Spearman’s rank correlation analysis was performed. Correlations were assessed among questionnaire items and task-performance scores (Table 1).

**Table 1.**
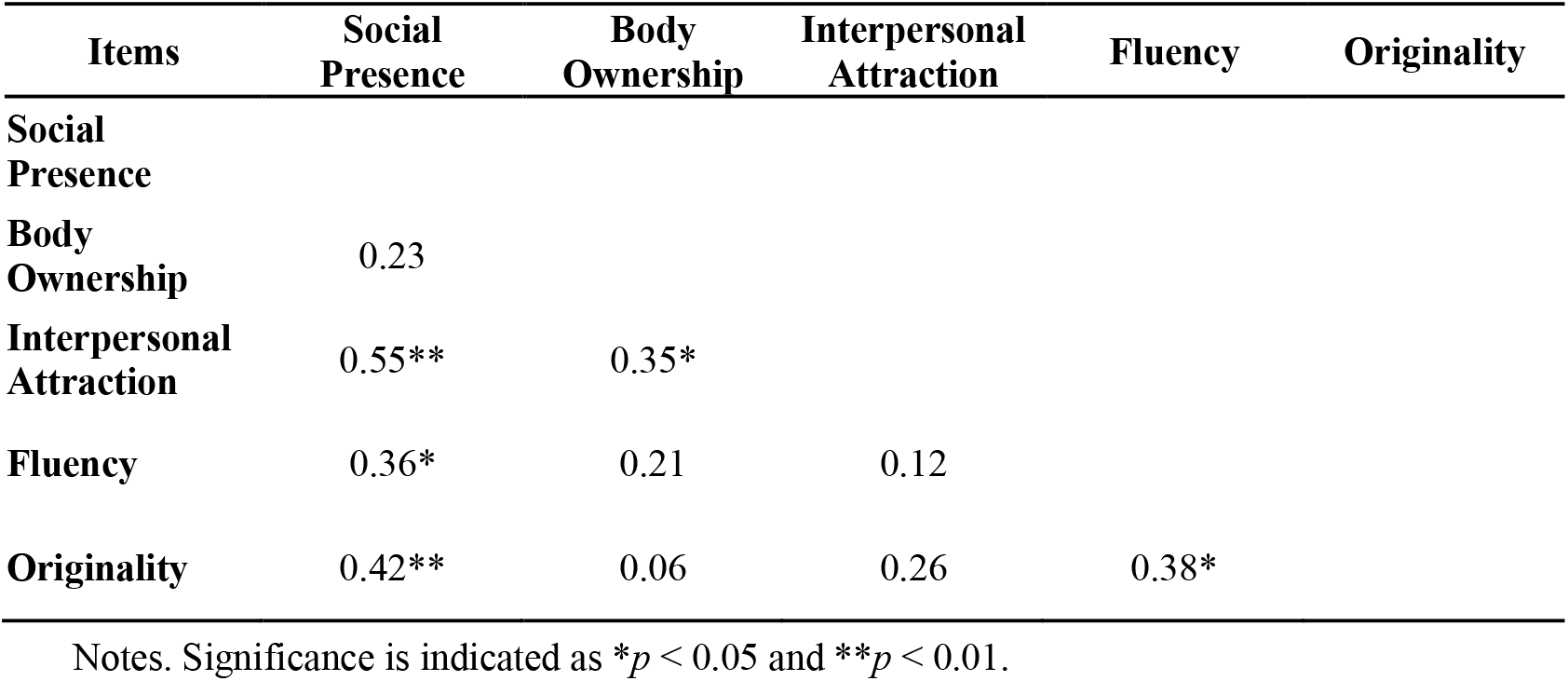
Spearman’s correlation matrix (*r* values) between subjective measures (Social Presence, Body Ownership, Interpersonal Attraction), task performance (Fluency, Originality).

| Items | Social Presence | Body Ownership | Interpersonal Attraction | Fluency | Originality |
| --- | --- | --- | --- | --- | --- |
| **Social Presence** |  |  |  |  |  |
| **Body Ownership** | 0.23 |  |  |  |  |
| **Interpersonal Attraction** | 0.55** | 0.35* |  |  |  |
| **Fluency** | 0.36* | 0.21 | 0.12 |  |  |
| **Originality** | 0.42** | 0.06 | 0.26 | 0.38* |  |

Notes. Significance is indicated as $*p < 0.05$ and $**p < 0.01$ .

Several significant positive correlations were observed: Social Presence–Interpersonal Attraction (*r* = 0.55, *p* < 0.01), Social Presence–Fluency Score (*r* = 0.36, *p* < 0.05), Social Presence–Originality Score (*r* = 0.42, *p* < 0.01), Body Ownership–Interpersonal Attraction (*r* = 0.35, *p* < 0.05), and Fluency Score–Originality Score (*r* = 0.38, *p* < 0.05).

These results reveal three notable patterns. First, higher levels of social presence and body ownership were associated with greater interpersonal attraction, suggesting that enhanced embodied and social experiences in VR are linked to stronger perceived social bonds. Second, social presence was positively associated with creative performance, as reflected in both fluency and originality scores on the Alternative Uses Task. This finding is consistent with prior work highlighting the role of social engagement in facilitating creative problem solving (Harvey & Kou, 2018). Finally, the positive association between fluency and originality supports previous evidence that these dimensions of creativity are interrelated.

## 4. Discussion

This study investigated how avatar facial expressions influence the interpersonal experience in VR by modulating the expressivity of the avatars used by participants. We employed a combination of subjective questionnaires, fNIRS-based hyperscanning, and task-performance comparisons. The questionnaire results revealed that facial expressions on avatars elicited a greater sense of body ownership, enhanced social presence, and increased interpersonal attraction. Furthermore, fNIRS data analysis showed that when avatars exhibited facial expressions, neural activity in the dlPFC and right TPJ—both of which are associated with social cognition—became synchronized between interacting participants.

The questionnaire results revealed a consistent pattern across all subjective measures, highlighting the central role of facial expressivity in shaping social experience within VR. Although task performance (fluency and originality) did not differ across conditions, the Amplified condition significantly enhanced both body ownership and social presence compared with the None condition.

Even though participants had only a limited view of their own virtual bodies, the visibility of their partner’s facial expressions appeared to increase their overall sense of “being there,” or presence, which in turn strengthened their sense of owning the avatar’s body. In this context, presence represents an individualized, context-dependent feeling of immersion or “being there” in the virtual environment (Bowman & McMahan, 2007). Social presence, as the interpersonal dimension of presence, was also highest in the Amplified condition, suggesting that the clearer an avatar’s facial expressions are conveyed, the stronger the sense of mutual “being together.” These findings imply a reciprocal relationship between embodiment and co-presence: when users perceive others as socially real, they also experience themselves as more embodied within the shared virtual space.

This interpretation contrasts with previous studies that used robot-like avatars limited to simplified eye and mouth movements during dyadic cooperation tasks (Dubosc et al., 2021). Such minimalistic avatars may fail to convey the subtle expressive cues necessary for emotional attunement and interpersonal resonance. In contrast, the combination of a human-like avatar and real-time facial expressivity in the present study likely induced more naturalistic social interactions, thereby enhancing both body ownership and social presence.

Finally, both the Amplified and Natural conditions yielded significantly higher interpersonal attraction scores than the None condition, replicating earlier findings that expressive avatars promote empathy, rapport, and affiliative behavior in VR interactions (Dubosc et al., 2021; Oh Kruzic, 2020). Taken together, these results suggest that the dynamic modulation of avatar facial expressions serves as a powerful nonverbal mechanism that integrates self-related (embodiment) and other-related (social connectedness) aspects of virtual experience.

The fNIRS results revealed significant increases in inter-brain synchronization (IBS) within the dlPFC and the TPJ. This pattern is consistent with previous studies demonstrating that IBS serves as a neural correlate of mutual understanding and effective social interaction (Cheng et al., 2015; Cui et al., 2012; Dikker et al., 2014; Funane et al., 2011; Osaka, 2014; Tang et al., 2016). Increased IBS within the dlPFC and TPJ may therefore reflect enhanced alignment of social-cognitive processes between interaction partners, supporting more coordinated and cooperative engagement during avatar-mediated communication.

The dlPFC has been widely implicated in executive control, working memory, and higher-order social-cognitive processes such as mentalizing (Decety, 2010; Lu et al., 2019; Vartanian et al., 2014). Increased IBS in this region during interaction may indicate a shared or mutually attuned cognitive stance between partners, reflecting reciprocal attention to each other’s intentions and task-related strategies (Xue et al., 2018). In the context of the present collaborative creativity task, which relies heavily on executive functions and working memory to generate and maintain multiple alternative ideas (Vartanian et al., 2014), enhanced avatar facial expressivity may have supported more effective engagement of these cognitive resources. Rather than directly improving task performance, such neural alignment may have facilitated a social-cognitive context conducive to sustained cooperation and joint problem solving.

Additionally, the middle frontal gyrus has been shown to be selectively engaged during the perception of dynamic facial expressions (Kessler et al., 2011) and to play a role in the cognitive control and regulation of others’ emotional responses (Vrticka et al., 2013). From this perspective, the observed increase in inter-brain synchrony (IBS) within the middle frontal gyrus may reflect participants’ active engagement in perceiving, monitoring, and interpreting each other’s facial expressions during interaction. Such neural coupling may index a shared attentional and regulatory process that supports the integration of expressive social cues in avatar-mediated communication.

The TPJ is widely recognized as a key hub for social cognition, particularly in mentalizing—the ability to infer others’ beliefs, intentions, and emotions (Saxe & Powell, 2006)—and in mediating joint attention during interpersonal coordination (Redcay, 2010). In our study, enhanced IBS in the right TPJ suggests that avatar facial expressions facilitated perspective-taking, encouraging participants to adopt their partner’s viewpoint. Such perspective alignment is a fundamental component of theory of mind, which supports mutual understanding and has been linked to improved team creativity (Lu et al., 2019). In this sense, avatar facial expressivity may have functioned as a catalyst for shared mental models, thereby strengthening the interpersonal resonance underlying creative collaboration.

Supporting this interpretation, Dravida et al. (2020) reported that TPJ-based IBS during face-to-face interaction is driven by reciprocal nonverbal behaviors, such as eye contact. It is therefore plausible that the inclusion of facial expressions in avatars enhanced mutual gaze between participants. Eye contact is known to foster reciprocal perception and emotional attunement in communication (Hirsh et al., 2017; Kaiser et al., 2022), and previous work has shown that mutual gaze increases social presence (Bente et al., 2008). Thus, the present findings extend this evidence to virtual settings, suggesting that expressive avatars can reproduce key elements of real-world nonverbal coordination that sustain social connectedness in VR.

Although these neural changes were evident, task performance (fluency and originality) did not significantly differ between conditions. This null result may stem from two sources. First, interaction style varied widely across pairs—some focused on idea generation, others on feedback or encouragement, and some exhibited prolonged silence—making quantitative comparisons difficult. Second, the open-ended nature of the Alternative Uses Task (AUT) allows for diverse response strategies, meaning that performance may have been influenced by factors unrelated to the experimental manipulation.

Nevertheless, the positive correlation between social presence and task performance indicates that a stronger sense of co-presence supports more effective creative collaboration. Previous studies have identified behavioral realism—including motion synchrony, gaze, and timely facial feedback—as a major predictor of social presence (Pütten, 2010). Our findings suggest that enhancing such realism through expressive avatars may, in turn, facilitate creative performance in VR-based collaborative contexts.

Despite these contributions, several limitations should be acknowledged. First, the participant sample consisted exclusively of young adults, which may limit the generalizability of the findings. Future research should include more diverse populations to examine potential age-related and cultural differences in avatar-mediated social interaction. Second, the present study focused solely on facial expressions as a nonverbal cue. Incorporating additional modalities, such as gaze behavior, body gestures, or vocal prosody, would provide a more comprehensive account of how multiple communicative signals jointly contribute to social presence and interpersonal coordination in VR.

Third, the collaborative creativity task permitted substantial variability in interaction strategies across pairs. Employing more structured collaborative paradigms, such as joint problem-solving or negotiation tasks, may help to isolate specific communication mechanisms and clarify how distinct social cues influence coordinated behavior.

## 5. Conclusion

This study examined how dynamically mapped and modulated avatar facial expressions influence body ownership, social presence, interpersonal attraction, and neural synchronization during VR interactions. Integrating subjective reports with fNIRS hyperscanning, we found that avatar facial expressions function as powerful nonverbal cues that strengthen the sense of presence and psychological connectedness in virtual environments. Notably, increased inter-brain synchronization within the dlPFC and right TPJ—both are associated with social cognition— suggests enhanced alignment of social-cognitive processes, potentially supporting perspective-taking and mutual understanding between interacting partners. This indicates that avatar facial expressions support deeper social cognition and mutual understanding. Although task performance (creativity scores) did not differ significantly across conditions, we found a significant positive correlation between creativity and social presence. This finding suggests that while performance may have been influenced by the task’s open-ended nature and variable interaction styles, social presence nonetheless plays a vital role in supporting creative collaboration. Overall, our findings demonstrate that expressive avatars are not merely visual features but are critical components for enhancing the quality of interpersonal interaction in VR environments. Leveraging avatar facial expressions to promote social cognition and emotional connection holds considerable promise for applications in communication and collaboration in virtual environments.

## Glossary

ANOVA: Analysis of variance
AUT: Alternative Uses Task
dlPFC: Dorsolateral prefrontal cortex
FDR: False discovery rate
IBS: Inter-brain synchrony
MNI: Montreal Neurological Institute
TPJ: Temporoparietal junction
VR: Virtual reality

## Acknowledgement

This research was supported by JST Moonshot R&D (Grant Number JPMJMS2013, Japan).

## Notes

### Competing Interest Statement

The authors have declared no competing interest.

